# Genome scan identifies flowering-independent effects of barley HsDry2.2 locus on yield traits under water deficit

**DOI:** 10.1101/216002

**Authors:** Lianne Merchuk-Ovnat, Roi Silberman, Efrat Laiba, Andreas Maurer, Klaus Pillen, Adi Faigenboim, Eyal Fridman

**Author notes:** These authors equally contributed to this manuscript (Lianne Merchuk-Ovnat, Roi Silberman). Author for correspondence: Eyal Fridman Institute of Plant Sciences, Agricultural Research Organization (ARO), The Volcani Center, P. O. Box 6, 5025001, Bet Dagan, Israel.

## Abstract

Increasing crop productivity under climate change requires the identification, selection and utilization of novel alleles for breeding. We analyzed the genotype and field phenotype of the barley HEB-25 multi-parent mapping population under well-watered and water-limited (WW and WL) environments for two years. A genome-wide association study (GWAS) for genotype by-environment interactions was performed for ten traits including flowering time (HEA), plant grain yield (PGY). Comparison of the GWAS for traits per-se to that for QTL-by-environment interactions (QxE), indicates the prevalence of QxE mostly for reproductive traits. One QxE locus on chromosome 2, *Hordeum spontaneum* Dry2.2 (HsDry2.2), showed a positive and conditional effect on PGY and grain number (GN). The wild allele significantly reduced HEA, however this earliness was not conditioned by water deficit. Furthermore, BC_2_F_1_ lines segregating for the HsDry2.2 showed the wild allele confers an advantage over the cultivated in PGY, GN and harvest index as well as modified shoot morphology, longer grain filling period and reduced senescence (only under drought), therefore suggesting adaptation mechanism against water deficit other than escape. This study highlights the value of evaluating wild relatives in search of novel alleles and clues to resilience mechanism underlying crop adaptation to abiotic stress.

**Highlight:** A flowering-time independent reproductive advantage of wild over cultivated allele under drought identified in a barley GWAS for genotype-by-environment interactions, with modified shoot morphology, reduced senescence and longer grain filling

## Introduction

Barley (*Hordeum vulgare ssp. vulgare* **L**.) is ranked the fourth most-produced cereal worldwide, providing fodder, human food and substrate for malting (Druka et al.,, 2011). Changing climate and increasing aridity poses a threat to future global food security (Wheeler et al., 2013), with drought stress, the major factor limiting crop yield (Boyer, 1982; Araus et al., 2008), expected to further increase (Wheeler and von Braun, 2013). It is therefore necessary to invest efforts in breeding-based improvements of crop resilience to abiotic stresses and in enhancement of plant yield robustness across a range of environments (Cattivelli et al., 2008; Tester and Langridge, 2010). Domestication of barley, approximately 10,500 years ago (Mascher et al., 2016), and its subsequent genetic selection, led to gene erosion (Tanksley and Nelson, 1996). Wild relatives of crops, which harbor most of the predomestication gene pool, may serve as valuable sources in attempts to formulate new genetic variation to improve drought resistance in modern varieties (Ellis et al., 2000; Zamir, 2001).

Traits enabling drought tolerance may involve adaptive phenological and cellular processes responsive to water stress. This includes 'escape mechanism', by an extremely early flowering at the expanse of shorten growth cycle; 'drought avoidance' in which active accumulation of solutes (which reduces osmotic potential to a more negative value) retain water in the cells (i.e., osmotic adjustment) and sustain metabolic activity (Blum, 2005). Plant survival only requires conserved water status and that is usually accompanied by growth inhibition. However, yield stability also requires the maintenance or increase in sink activity in the reproductive structures, which contributes to the transport of assimilates from the source leaves and to delayed stress-induced leaf senescence (Albacete et al., 2014). An ample number of genes have been suggested to be involved in drought tolerance, yet many were discovered using differential genomics methods (Bedada et al., 2014; Rollins et al., 2013). That ignore a whole-plant perspective, which is to key to understanding the subtle sink-source relationship and its optimal maintenance for yield stability. Genome scans in search of reproducible quantitative trait loci (QTL) by-environment interaction (QxE) loci under field set-ups, which takes into account possible pleiotropic effects, or lack thereof, provide a promising entry point for deciphering major drivers of pathways that confer field drought tolerance.

Advanced backcross QTL analysis (AB-QTL) (Tanksley and Nelson, 1996) enables the introduction of beneficial alleles into the modern gene pool, by crossing a wild donor accession with a modern elite cultivar (Fulton et al., 2000; Pillen et al., 2003; Talamè et al., 2004; Von Korff et al., 2005; Von Korff et al., 2006; Von Korff et al., 2008; von Korff et al., 2010), followed by a number of selfings. Also, recombinant inbred line (RIL) populations, derived from crossings of a wild donor with a cultivar (or between two distinct domesticated parental lines), have been used to map QTLs for grain yield and yield components under reduced moisture (Teulat et al., 2003; Kirigwi et al., 2007). In contrast to the limited allelic diversity in such bi-parental-based genetic structures (Comadran et al., 2011), the recently developed multiparental population, that combines linkage and genome-wide association (GWA) approaches, offers a much wider genetic variance. GWA studies (GWAS) have seldom been used to detect QxE interactions, mainly due to lack of statistical power, owing to the frequent occurrence of rare alleles that are difficult to detect in a GWAS, but which appear to contribute to strong genotype by-environment interactions (Thomas, 2010). Notably, in most of these studies, the genetic model undertaken compares the GWAS results under one versus another environment to identify environment-specific QTLs. Very few studies have considered marker-by-environment interactions in their genetic model, and thereby preclude testing of same SNPs across all environments and reduce the ability to discover novel alleles or genes that synergistically contribute to environmental adaptation and plasticity (El-Soda et al., 2014). From a breeding point of view, constitutive QTL are the main targets for breeding programs, as they show a consistent effect across environments (Comadran et al., 2011; Korte et al., 2012). However, if the goal is to understand the mechanism underlying GxE interactions, then conditioned QTL are imperative targets for follow-up studies.

The barley nested association mapping (NAM) population, termed 'Halle Exotic Barley 25' (HEB-25), originated from interspecific crosses between the spring barley elite cultivar Barke (*Hordeum vulgare* ssp. *vulgare*, Hv) and 25 highly divergent exotic *H. spontaneum* (Hs) wild barley. The population was used to study the genetic architecture of flowering (Maurer et al., 2015) and grain weight (Maurer et al., 2016). The NAM approach (Buckler et al., 2009) was originally designed to overcome GWAS limitations, such as the incidence of false positives resulting from population structure. In this study, NAM was harnessed to evaluate marker by-environment interactions. Each of the 1420 HEB-25 BC_1_S_3_ lines and their corresponding parents were genotyped using the Infinium iSelect 9K chip, which consists of 7864 SNPs (Comadran et al., 2012). Previously, the combined linkage and GWAS analysis of HEB-25 identified eight major QTL that control flowering time, potentially explaining the QTL effect. Most co-located with major flowering genes, including Ppd-H1, HvCEN (2H) and Vrn-H1/H2/H3 (on chromosome 5H, 4H and 7H, respectively). The strongest exotic haplotype identified accelerated flowering time by 11 days, as compared to Barke (Maurer et al., 2015). Similarly, Maurer et al. (2016) reported that grain weight was increased by 4.5g and flowering time was reduced by 9.3 days after substituting Barke elite QTL alleles for exotic QTL alleles at the semi-dwarf locus *denso/sdw1* (3H) and the *Ppd-H1* loci, respectively. In a more recent use of this genetic resource, the plants were placed under two regimes of salinity and the GWA of the two treatments was compared (Saade et al. 2016). While marker by-environment interactions were not reported, constitutive QTL under both environments, which, from a breeding point of view is of high value, were identified.

In the current study, the BC_1_S_3_ HEB-25 families were used in a GWAS of the genetic architecture of drought response, as defined by plant grain yield and related traits. A mixed linear model tested the QxE for the studied traits in order to identify specific loci that contribute to plant adaptation via dependent or independent effects on phenology and morphology. The analysis showed no interactions between flowering time loci and identified several significant interactive QTL that affect plant grain yield and number in a water deficiency dependent manner. Furthermore, a pot experiment was carried out using an advanced backcross population segregating for the HsDry2.2 locus which was found to improve yield when donated by the wild parent at this locus. This experiment was designed to evaluate the effect of an early and late water limitation, by measuring plant productivity, phenology, canopy structure and leaf dimensions. This study highlighted the power of integrating semi-controlled field experimental systems and inter-specific multi-parental populations in pursuit of agricultural traits, which may shed light on hitherto unknown mechanisms underlying crop adaptation.

## Materials and methods

### Plant Material

#### Field trials

The NAM population HEB-25, which was developed by Maurer *et al.* (2015), consisted of 1,420 BC_1_S_4_ lines belonging to 25 interspecific crosses between cv Barke and wild barley accessions. All 1,420 BC_1_S_3_ lines (one generation earlier) and their corresponding parents were genotyped using the barley Infinium iSelect 9K chip (Maurer et al., 2015), consisting of 7864 SNPs (Comadran et al., 2012). Inclusion of SNPs that were polymorphic in at least one HEB family and that met the predefined quality criteria (<10% missing, and not in complete linkage disequilibrium (LD) to another SNP in the set) resulted with 5709 informative loci.

#### Pot experiment

BC_2_F_1_ seeds (HEB-04-96) segregating for the wild donor allele in the HsDry2.2 locus were genotyped for the peak marker BOPA2_12_30265 using high resolution melting analysis. This marker was formerly found to show strong divergent selection in winter *vs*. spring barleys according to Comerdan et al., (2012). SYTO™ 9 Green Fluorescent Nucleic Acid Stain (ThermoFischer, 0.6µl per 20µ reaction) was used for PCR along with Taq ready mix (HyLabs), with the primers listed in Table S1. Melting analysis was conducted using RotorGene 6000 real-time PCR machine and software (Fig. S1A.) and was the validate by sequencing (Fig. S1B). Barke cultivar, which sets the genetic background was also included and genotyped as control. The different genotypes were grouped as carriers for the wild (Hs/Hs and Hv/Hs; N=22 and 7, respectively) or cultivated (26 Hv/Hv; N=26) allele for statistical analyses.

### Field trials growth Conditions

The HEB-25 lines were evaluated for their drought responses under well-watered (WW) and water-limited (WL) conditions during the winters of 2014–15 (BC_1_S_3:4_) and 2015–16 (BC_1_S_3:5_). During 2014-15 the entire population was phenotyped in the field under both conditions, and in 2015-16 1320 lines were included in the experiment. Sixteen (2 groups of 8) plants from each line were randomly arranged in an insect-proof screen house, roofed by polyethylene, at the experimental farm of The Hebrew University of Jerusalem in Rehovot, Israel (E34°47′, N31°54′; 54m above sea level). Plants were grown within paired troughs (Mapal Horticulture Trough System, Merom Golan, Israel) that allowed regulation of watering throughout growth (Fig. 1A and 1B). The unique arrangement of the nethouse enables examining app. 3000 experimental units of 8 plants, with capacity to split irrigation between adjacent troughs. Three cultivated barley lines (Apex, Barke and Bowman) served as control for the effects of the water deficit. The soil in the troughs was composed of two layers of volcanic soil (4-20 topped by Odem193 (Toof Merom Golan, Merom Golan, Israel)). The size of mini-plots, each containing eight plants, was 0.4 × 0.3 meter (Fig. S2A). To mimic the natural pattern of rainfall in the east Mediterranean region, water was applied during the winter months starting from planting (Dec. 14 2014 and Nov. 23 2015, respectively) and ending in early spring (143 and 145 days after planting, respectively) (Fig. S2B). In order to ensure adequate drought stress, irrigation was adjusted in accordance with the stomatal conductance determined in eight test plants per irrigation treatment during the growing season (57, 98, 110, 127 days from planting in 2014-15, and 70, 84, 119, 154 days from planting in 2015-16), measured using a porometer (Decagon SC-1) (Fig. S2C). This assured a ~30% differences between the WL and the WW plants, with maximal conductance of 350 and 250 mmol m^−2^s^−1^ in the WL and WW treatments, respectively. The total seasonal water application included 25m^3^ and 34m^3^ for the WW treatment, and 13m^3^ and 26m^3^ for the WL treatment, in 2014-15 and 2015-16, respectively. NPK fertilizer (Shefer 538 + Microelements, Deshen Gat, Qiryat Gat) was applied via irrigation in 2014-15 (8.1 and 6.5 liter for the WW and the WL treatments, respectively) as well as in 2015-16 (15.4 for both treatments).

**Fig. 1.**
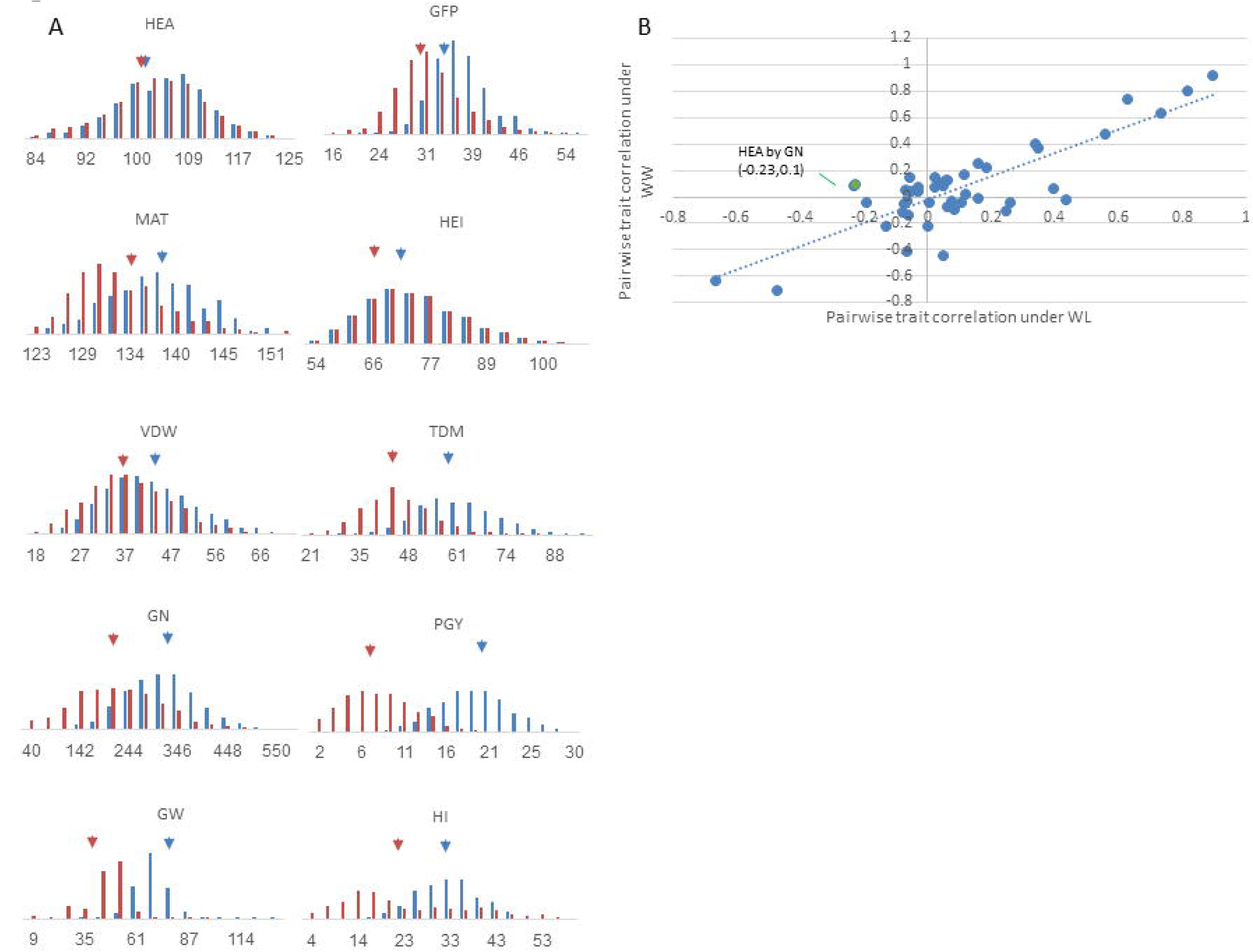
Distribution and correlation matrix of the measured traits under WW and WL conditions. (A) The distribution of the traits is presented for the HEB-25 lines for the WW (blue) and WL (orange) irrigation conditions (see detailed values in Table S1). Blue and orange arrows facing down depict the mean values of the HEB-25 population under WW and WL environments. (B) Scatter plot of the pairwise correlations between the 10 traits under WL (X-axis) and WW (Y-axis). HEA, days from sowing to anthesis; MAT, days from sowing to maturation; GFP, Grain filling period [days]; HEI, plant height [cm]; VDW, vegetative dry matter [gr]; TDM, total dry matter [gr]; PGY, plant grain yield [gr]; GN, grain number [#]; GW, grain weight [mg].

### Pot experiment irrigation management

Seeds were placed in moist germination paper for a week in a dark cold room (4^°^C), followed by a three days of acclimation at room temperature (22^°^C), then planted into small plastic pots (60g soil and 300g water, 100% field capacity) in a standard commercial soil mix (Green 90, Even Ari, Israel) (Fig. S2D). At 28 days after planting (DAP) plants were transplanted into medium pots (190g soil and 1170g water). Finally at 52 DAP plants were into large pots (460g soil and 2500g water), with the initiation of the late drought treatment, 13 days before booting (Fig. S2E). Pots were weighed manually before and after irrigation, keeping the well -watered (WW) pots between around 60-90% of field capacity and the water limited pots (WL) 40-60%. Temperatures were monitored using a data logger (Hobo Onset, Bourne, MA USA).

### Phenotypic measurements

#### Field trials

Heading time (HEA), defined as the time between sowing to time at which the first spike of 50% of the plants in a plot reaches BBCH49 (first awns visible), was recorded based on daily inspection. Days from sowing to stage BBCH87 (hard dough: grain content solid: fingernail impression held) was recorded as maturity (MAT). At maturity, plant height (HEI) was measured from the soil surface to the base of the three first spikes per plot.

At full grain maturity and after plants were fully dried, all aboveground biomass was harvested and weighed to determine total dry matter (TDM). Notably, all the free-thrashing material (app. ¼ of the material) was caged between BBCH49 and BBCH87 to avoid loss of spikes. Spikes were then threshed and weighed to determine plant grain yield (PGY). Finally, grains were counted to estimate grain number per mini-plot (GN) and average grain weight (GW). Harvest index (HI) was calculated as the ratio between PGY and TDM. Vegetative dry matter (VDW) was calculated by subtracting PGY from TDM. The grain-filling period (GFP) was calculated by subtracting HEA from MAT. Trait values were adjusted based on the ratio between population mean values in the two years. The adjusted HEB means across years were used in the GWAS.

#### Pot experiment

Booting time, defined as the date at which the three spikes awns are first visible in a pot (spikes are tagged), was daily recorded and used to score HEA. MAT was recorded when the three first (tagged) spikes dried, which was right followed by the rest of the plant drying out. GFP was calculated by subtracting HEA from MAT. All aboveground biomass per pot was harvested at full grain maturity; fertile spikes were counted to assess the number of spikes per plant, separated from the vegetative organs (stems and leaves), and both were oven-dried (80^°^C or 38^°^C for 48 h for vegetative organs and spikes, respectively) and weighed to determine spike dry matter and TDM. Spikes were threshed and total grain weight was determined. Plant grain yield (PGY), harvest index (HI = PGY/TDM), grain number per pot (GN) and averaged grain weight (GW) for the three first headed spikes (tagged) and the rest of the spikes were calculated separately. At 64 DAP, FL sheath and ‘minus one’ blade length and blade width were scored. Stem diameter (adjacent to the spike) was scored after harvest. For the analyses of leaf senescence we used two sets of photos took using Canon EOS1200 camera at 66 and 78DAP in the background of standard white sheet (80×120 cm). A custom-made software (unpublished) was used to calculate the ratio of the yellow-brown shades out of the total leaf area. Shades that cover the range of green or yellow brown were selected by sampling from several images, and were then used for all analyses. For each photo, the green area and the yellow-brown area were calculated in percentages, using the YCbCr method, with same tolerance level for al analyses, while taking into account only the white background. Then, the percentage of yellow-brown out of total leaf (yellow brown+green) area was calculated. Senescence was calculated as the delta of yellow-brown/total between dates (78-66 DAP) for each pot.

### Field trails genome-wide association study (GWAS)

A genome-wide association study was conducted to identify trait variations, per-se, under WW and WL conditions, and to assess SNP interactions with the environment (conditional effects), using the GWAF package (<u>http://cran.r-project.org/web/packages/GWAF/</u>). To test association between each of the continuous traits and each SNP, we applied the linear mixed effects model (LME) implemented in lmekin function in kinship package (<u>http://cran.r-project.org/web/packages/kinship/</u>). The analysis was conducted with the adjusted means following a linear mixed model as follows:

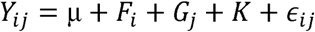

where, μ denote the population mean for the trait, F_i_ is the family effect (*i*=1..25), G_j_ denotes the marker effect (including heterozygous, i.e. j=1..3), K represents the relative kinship matrix, and ∈_ij_ denotes the error.

A genome scan for SNP x environment (QxE) interaction was conducted with the GWAF package, using the same LME model, but with addition of the WW/WL condition as follows:

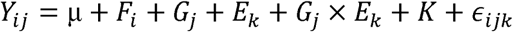

where, *E*_*k*_ denotes the watering (environment) effect, *G_j_* X *E_k_* denotes the effect of the interaction between the marker and the environment, K represents the relative kinship matrix, and *∈_ijk_* denotes the error.

To test the robustness of the association for each SNP, the procedure was cross validated 200 times on random sub-samples of the full dataset. Each subsample included 70% of the lines, randomly selected per HEB family. Markers that were significantly detected (P<0.05) in at least 30% of sub-samples were accepted as putative QTLs. The designation of the QTL was based on LD with the major SNP that showed the maximal significance level in a genomic interval (Maurer et al. 2015).

### Pot experiment statistics

A factorial model (3 irrigation treatments x 2 allelic states) was employed for the analysis of variance (ANOVA), with irrigation treatment and allelic state as fixed effects. Each experimental unit consisted of a pot with one plant. The JMP version 12.0 statistical package (SAS Institute, Cary, NC, USA) was used for statistical analyses.

## Results

### Whole-plant phenotype of the HEB-25 multi-parent population under well-watered (WW) and water-limited (WL) conditions

The HEB-25 family was grown in mini-plots of 8 plants during 2014-15 and 2015-2016 (hereafter, 2015 and 2016). The experimental set-up for the mini-plots under a rain sheltered nethouse (high-content plant phenotyping; HCPP) was designed to allow whole-plant phenotype of plants, and at the same time to allow controlled irrigation with validated water deficit in the plants (Fig. S2A). Fig. S2B depicts the accumulated irrigation throughout the experiment during 2016. Assessment of the effect of WL on stomatal conductance of control plants that were distributed across the entire HCPP system, clearly indicated mild, yet statistically significant differences in the stomatal conductance between WW and WL starting at least 70 days after sowing (no porometer measurements were made in younger plants), and onward (Fig. S2C).

Overall, we observed a wide variation for all traits across HEB-25 and increase in the coefficient of variation under WL, i.e. mean of 28.3 vs 16.8 for all traits under WL vs WW, respectively (Fig. 1A, Table S2). WL conditions significantly reduced means of all measured traits, as compared to WW conditions, with effects on heading time (HEA) being the mildest (-1.2%). On average, plant grain yield (PGY) and grain number (GN) were reduced by −58.3% and −31%, respectively, under WL conditions, as compared to WW. Effects of WL on vegetative parts were weaker, with a −13% reduction in vegetative dry weight (VDW) and −13 % reduction in plant height (HEI). Pair-wise correlation analysis between all traits found similar relationships under both treatments, with few exceptions (Fig. 1B, Table S3). One such exception is the relationship between HEA and GN; under WW conditions there was a positive and mild correlation between these traits (r=0.1, P<0.005), whereas under WL, this correlation was negative (r=-0.23, P<0.0001).

### GWAS for QxE interactions

Initially, we performed a genome scan for all 10 traits, under both treatments. For the 10 traits examined in this study, 69 loci with a significant contribution to trait variation were identified (Fig. 2, Table S4). The main loci for HEA, including HvELF3, Ppd-H1, HvCEN, *denso*, Vrn-H1, and Vrn-H3, were identified. In general, at loci associated with PGY and GN the wild alleles were associated with a reduction of the traits under WW conditions. Under this trait per se analysis the mean reduction of the three loci significantly associated with GN or PGY variation, respectively, were −3.4% and −9.2% under WW conditions (Table S4). Similar effects were observed under WL conditions for a locus linked with Ppd-H1 on chromosome 2 (-19.7%; - LogP=7.9) for GN.

**Fig. 2.**
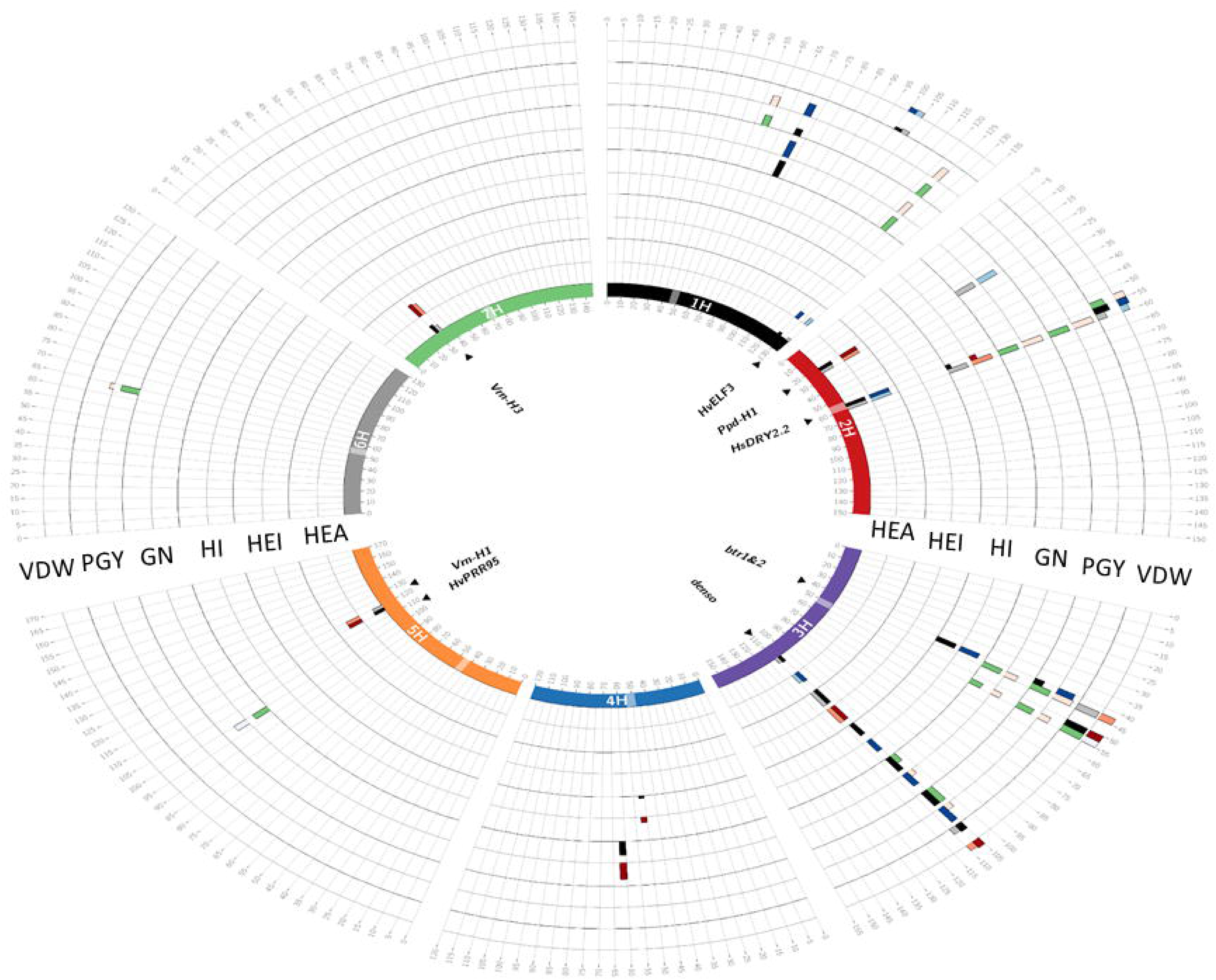
Circos plot depicting the GWAS results for trait per-se (QTL) and interaction with environment (QxE). Barley chromosomes in the Circos plot are depicted in different colors the inner circle and centromeres are indicated by transparent boxes. For each trait, the first (inner) track represents the –Log_10_P score of QTL detection in a 5-cM window and the adjacent outer track represents the effect of this QTL. The maximum height of the effect bars is 10.03 days for HEA, 17.34Lcm for HEI, 24.3 for HI, 22.5 for GN, 22.44 for PGY and 23.04L for VDW. Window positions (in cM, as per Maurer *et al*. 2015) are ordered clockwise, per chromosome. In the inner track, QTL appearing under WW and WL conditions are presented by black and gray bars, respectively. The QTLs showing significant QxE interactions are represented by green bar. The effect of the QTL conferred by the wild allele relative to Barke is represented on the outer track, where blue and red bars indicate decreasing and increasing wild barley QTL effects, respectively, for each treatment. Genes potentially explaining the observed QTL effects, are indicated inside the inner circle.

Next, the GWAS model was modified to identify QTL by-environment interactions (QxE; see Methods). No significant QxE loci were identified for two of the phenological traits HEA and MAT. For the vegetative traits, only two QxE loci were identified in VDW and no significant loci were identified for plant height (Table S5). In contrast, a larger number of interactive loci were associated with PGY or GN variation (Fig. 2). In most of these loci the wild allele was characterized with a conditional beneficial effect on the trait, i.e. carriers of the wild allele were less affected by the water deficit across the two environments.

### HsDry2.2, a major locus with non-pleiotropic QxE interactions

One major locus that appeared as a pleiotropic QTL that regulates many traits is located on the long arm of chromosome 2 and delimited by SCRI_RS_144592 and SCRI_RS_165574 (54.2-59.9 cM; Fig. 3). We named this locus *Hordeum spontaneum* Dry 2.2 (HsDry2.2). The peak of the QxE interaction in this locus was identified at 57cM. Plotting the mean values of PGY measured for the HEB-25 population under WW and WL conditions, illustrated the conditioned effect of HsDry2.2 on the trait (Fig. 4A). This is in contrast to identical effects under both growth conditions of the wild allele on HEA or VDW (no interaction; Fig. 4B; Fig. S3), i.e. reduction of the days to heading by an average of 7.7 days and 8.1 days in WW and WL, respectively. Conditioned effects on GN were more pronounced (Fig. 4C), and no significant effect on mean grain weight (GW; Fig. 4D) was associated with the locus.

**Fig. 3.**
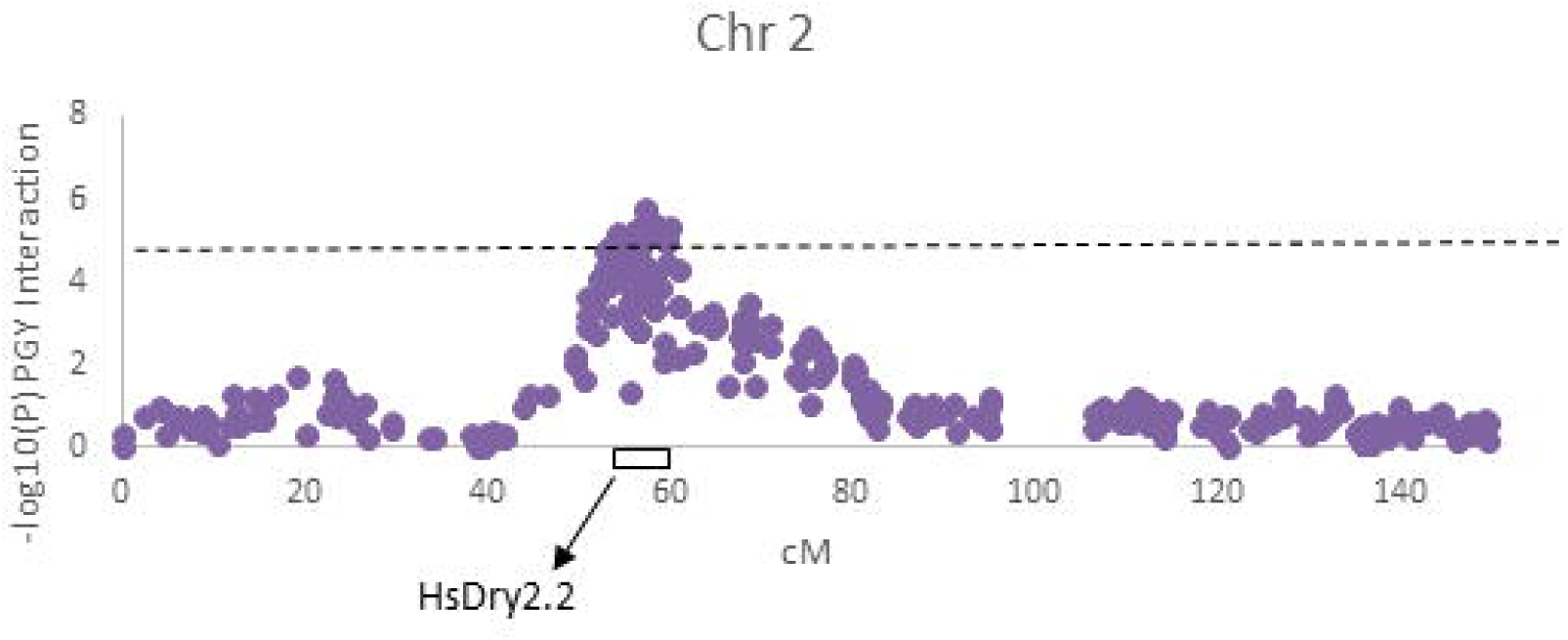
Genetic mapping of the genome wide association for QxE interactions for PGY in chromosome 2. Manhattan plot depicting the location of HsDry2.2 locus on the long arm according to genetic map. The Y axis depicts the –log(P) value of the QxE interaction. It is delimited by SCRI_RS_144592 and SCRI_RS_165574 (54.2-59.9 cM), peaking at 57cM.

**Fig. 4.**
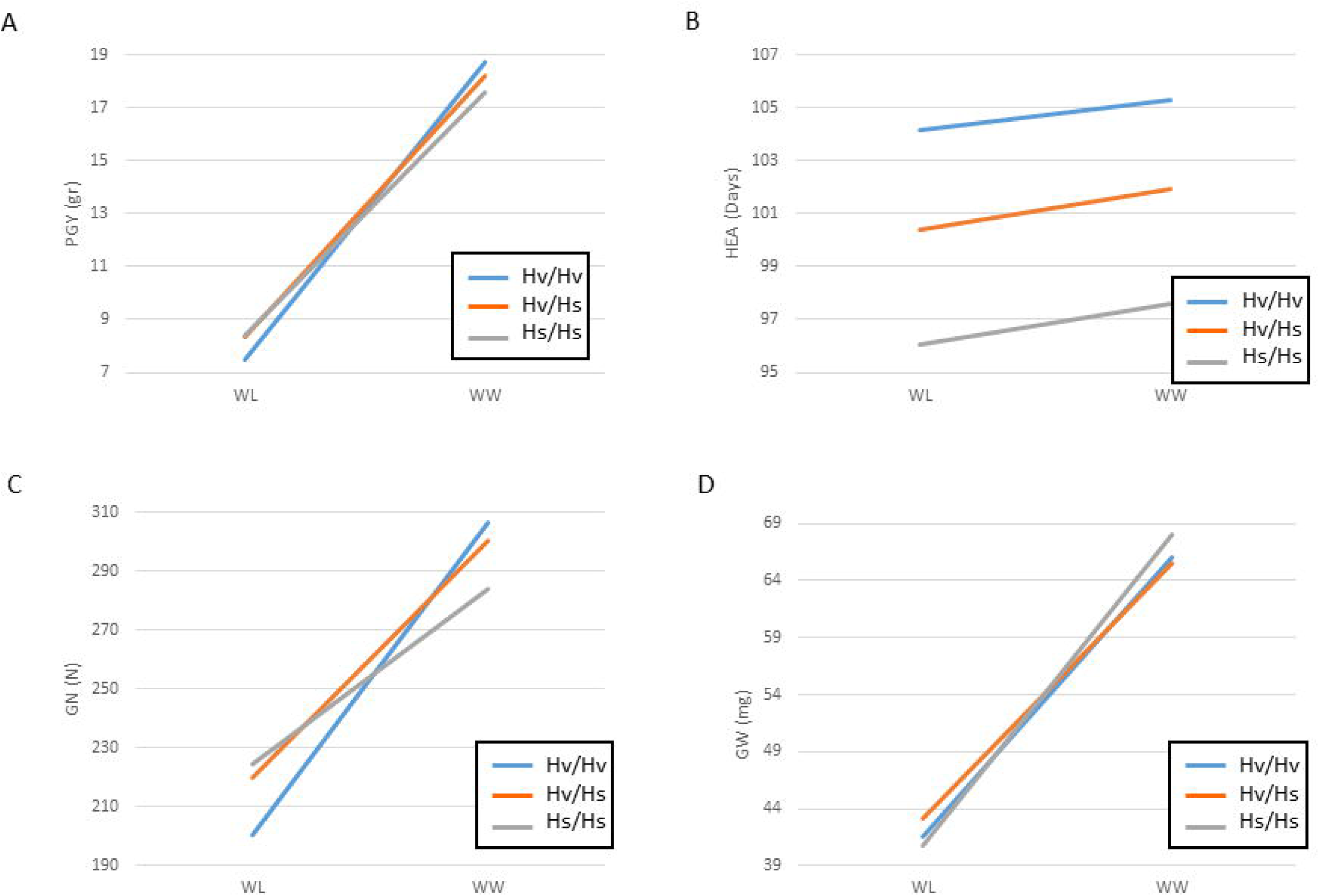
Least square mean value comparisons for the *HsDry2.2* genotypes under WW versus WL (reaction norms) conditions, in the whole HEB-25 population. Reaction norms for (A) plant grain yield (PGY), (B) days from sowing to heading (HEA), (C) grain number (GN), and (D) grain weight (GW). The three genotypic groups of plants homozygous for the Barke cultivated allele (Hv/Hv), homozygous for the wild allele (Hs/Hs) and heterozygous (Hv/Hs) are depicted by blue, ornage and gray lines, respectively.

Dissection of the family-specific effects of HsDry2.2 on GN, suggested that the causal allele or alleles for this significant QxE interaction (P<0.01) originated from at least two wild donor. Performing the genotype to phenotype analysis in the HEB-16 family shows that plants carrying the wild HID219 allele (Maurer et al. 2015) exhibited a significant reduction in GN between WW and WL (334 vs 221 in WW and WL, respectively, i.e., a reduction of 34% under water deficit) in a similar manner to that of the cultivated allele carriers (Fig. S4A). On the contrary, a much milder reduction in plants carrying the wild allele was observed in the HEB-5 family (HID065 as donor) with a reduction of 7.3% from 272 to 252 for the homozygous to the wild allele (Fig. S4B). These differences translated to a family-specific positive conditioning effect of the wild barley allele under WL conditions.

### The effect of HsDry2.2 in consecutive BC_2_S_1_ population

Plants derived from one of the HEB lines that carry the HsDry2.2 *H. spontaneum* (HID062) allele were further analyzed to validate effects on reproductive traits, as well on possible related traits. A pot experiment was carried out to evaluate the wild allele effects under optimal as well as early and late water limitation, i.e. starting at transplanting or at stem elongation stage (52 days after planting; 52DAP), respectively. Type of traits included plant productivity, phenology, canopy structure and leaf senescence.

#### Overall plant performance

Irrigation treatment effect was found significant for most of the measured variables (Fig. 5, Table S6). TDM under the Early WL and Late WL treatments was reduced by 31% and 26% as compare to the WW treatment, respectively. GY was reduced by 17% and 16% for the early and late WL as compare to the WW. HI increased under both WL treatments as compare to the WW (16% and 12% for the early and late respectively). Leaf blade parameters −1FL width and length also exhibited reduction of 10% and 3%, and 16% and 10% for the traits and treatments respectively. Whereas all of these mentioned above traits showed similar levels under both WL treatments, both GW and senescence showed significant differences between the early and late drought: GW increased with drought level by 17% and 9% for the early and late WL, and canopy senescence increased by 62% and 38% for the early and late respectively (Fig. 5). About 4 days earlier heading (smaller HEA, 6%) was recorded under the Early WL relative to the WW, whereas the Late WL did not differ from the WW as expected since Late WL initiated only at 52 DAP. The number of spike per plant was also reduced by the early but not by the late WL (14%). Irrigation treatment showed no significant effect on grain number (Fig. 5).

**Fig. 5.**
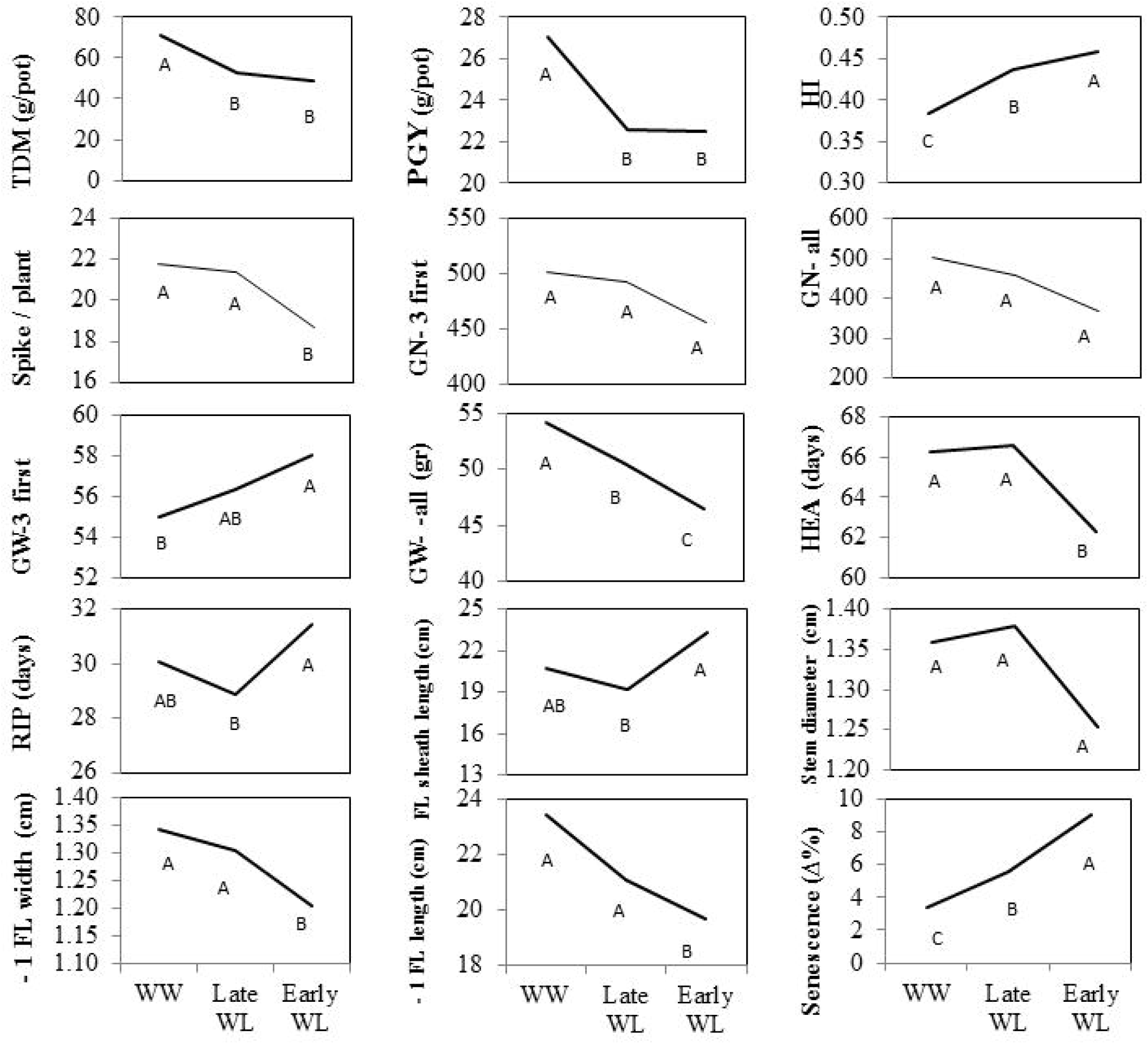
Effect of irrigation treatment on the different parameters evaluated: total DM (TDM, gr/plant), grain yield (PGY, gr/plant), harvest index (HI), spike per plant, grain number of three first spike (GN-first three), GN all, grain weight of three first three spike (GW-first three), GW, all, days from planting to booting (HEA, days) ripening period (RIP, days), flag leaf (FL) sheath length (cm), stem diameter (cm), −1 FL blade width (cm), −1 FL blade length, and senescence.

### Productivity related traits

After genotyping the BC_2_S_1_ plants for the three genotypic groups (Fig. S1), and observing dominant mode of inheritance for most traits (data not shown), we decided to group the carrier of one or two wild alleles as a single group (hereafter Hs/_) to increase the number or replicates. TDM did not differ between the Hs/_ and the cultivated (Hv/Hv) groups averaged across treatments and under each treatment separately (Fig. 6A). However, the wild allele conferred an advantage in PGY averaged across treatments (Fig. 6B). Under the WW treatment this advantage was found most pronounced. Harvest index of the wild allele carriers was significantly higher than in the cultivated allele group, especially under the Early WL (Fig. 6C). Spike number per plant did not significantly differ between genotypes under the different treatments, or averaged across treatments (Fig. 6D). The first three spikes to boot were weighed and threshed separately. Grain number (GN) of the first 3 spikes was significantly higher in the Hs/_ than in the Hv/Hv group (Fig. 6E). Genotype by treatment interaction curve (reaction norm) shows this effect was obtained under both the Late WL and the WW, but not under Early WL. GN of all spikes was higher (p=0.08) for the Hs/_, when averaged across treatments, and under the WW treatment (Fig. 6F). While the wild allele was associated with lower GW of the first three spikes as compare to the cultivated one (mainly affected by the WW, P=0.06, Fig. 6G), an opposite trend was observed for GW of all spikes (mainly having an increasing effect at the Early WL, P=0.07, Fig. 6H).

**Fig. 6.**
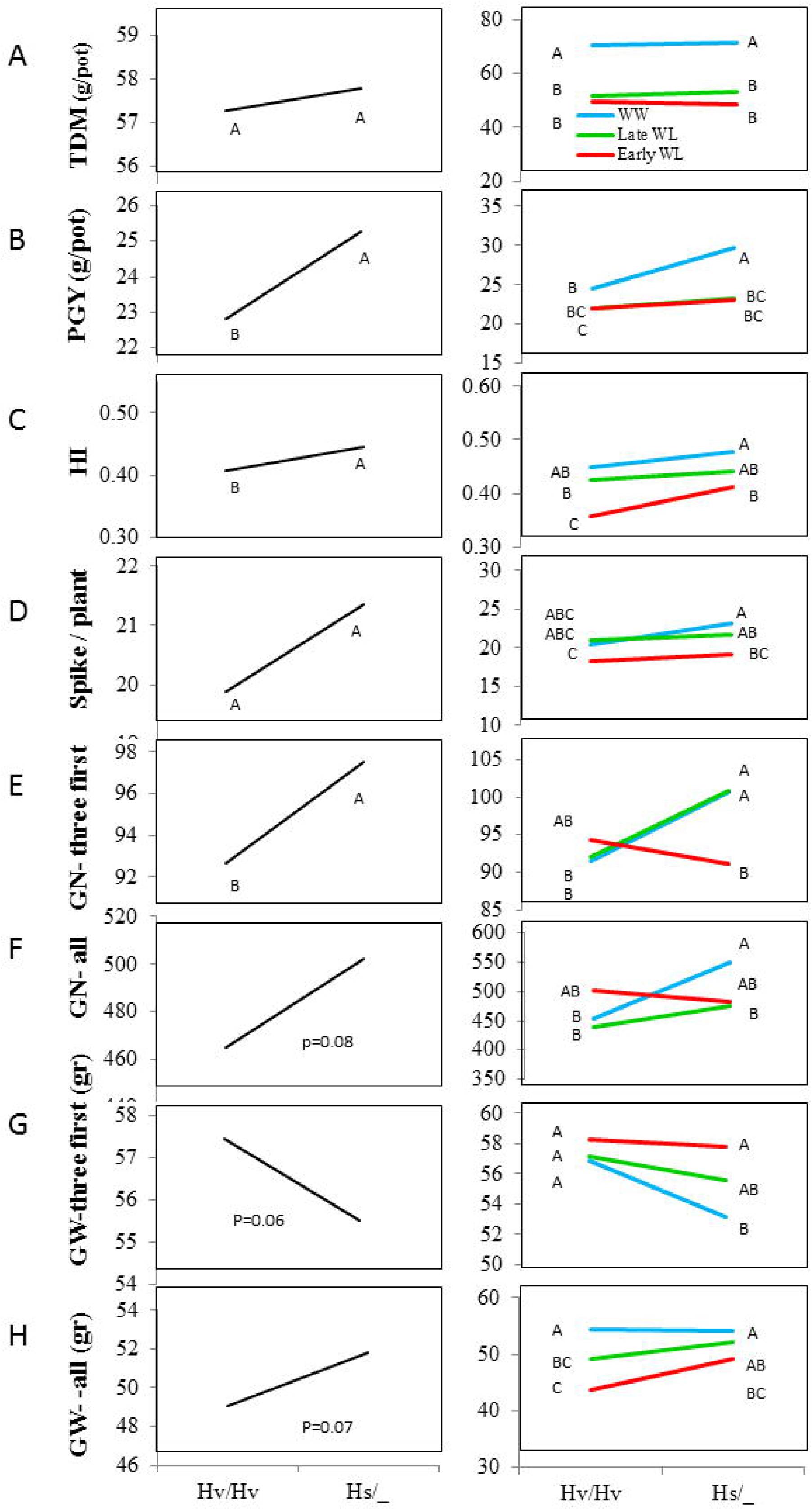
Genotype and genotype by environment interaction curves of HsDry2.2 for productivity related traits: (A) total DM (TDM, gr/plant), (B) plant grain yield (PGY, gr/plant), (C) harvest index (HI), (D) spike per plant, (E) grain number of three first spike (GN-first three), (F) grain number (GN all), (G) grain weight of three first spike (GW-first three), (H) grain weight (GW, all). Hv/Hv, homozygous for the Barke cultivated allele, Hs/_ homozygous or heterozygous for the wild HID062 allele.

#### Phenology

The first three spikes to head were used to determine plant phenology. The Hs/_ group booted significantly earlier than Hv/Hv plants (2.5 days Fig. 7A), which was more pronounced under WW and Late WW than in the Early WL. The wild allele carriers presented an elongated ripening period across all treatments as compare to the cultivated allele (2 days, Fig. 7B).

**Fig. 7.**
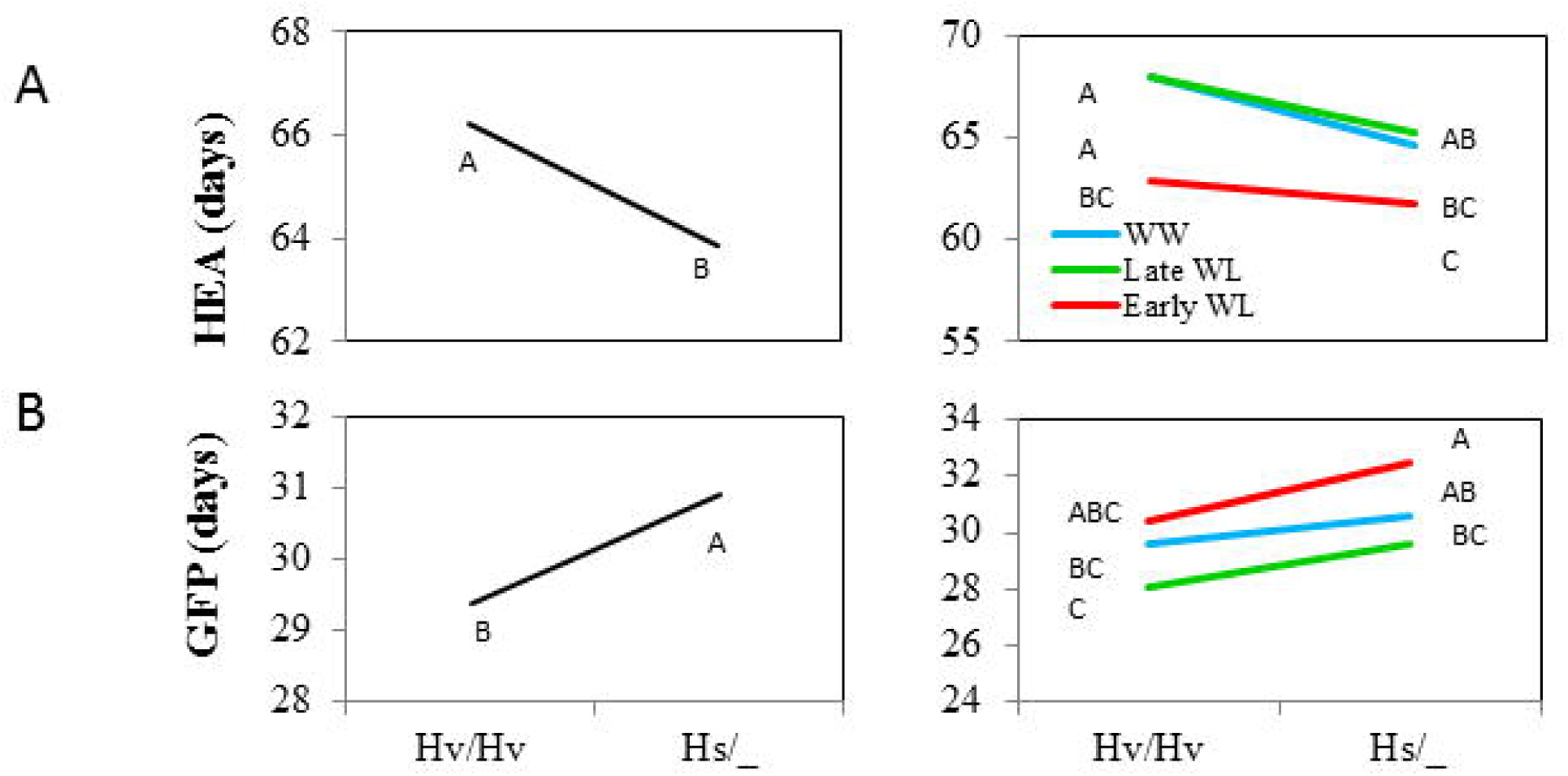
Genotype and genotype by environment interaction curves for phenological traits: (A) days from planting to booting (HEA, days), (B) ripening period from HEA to maturity (RIP, days). Hv/Hv, homozygous for the Barke cultivated allele, Hs/_ homozygous or heterozygous for the wild HID062 allele.

#### Canopy structure

Flag leaf sheath length was higher in the Hs/_ group (Fig. 8A and Fig. S5A). The wild allele effect was very stable in this parameter across treatments and the strongest differences between genotypes were observed under Late WL. The Hs/_ plants exhibited constitutively reduced stem diameter (Fig. 8B and Fig. S5B). The wild allele was generally associated with longer and narrower leaves; Both 'minus one' flag leaf blade width and length were found to be lower for wild allele carriers with most pronounced difference under Early WL (Fig. 8C and 8D; Fig. S5A).

**Fig. 8.**
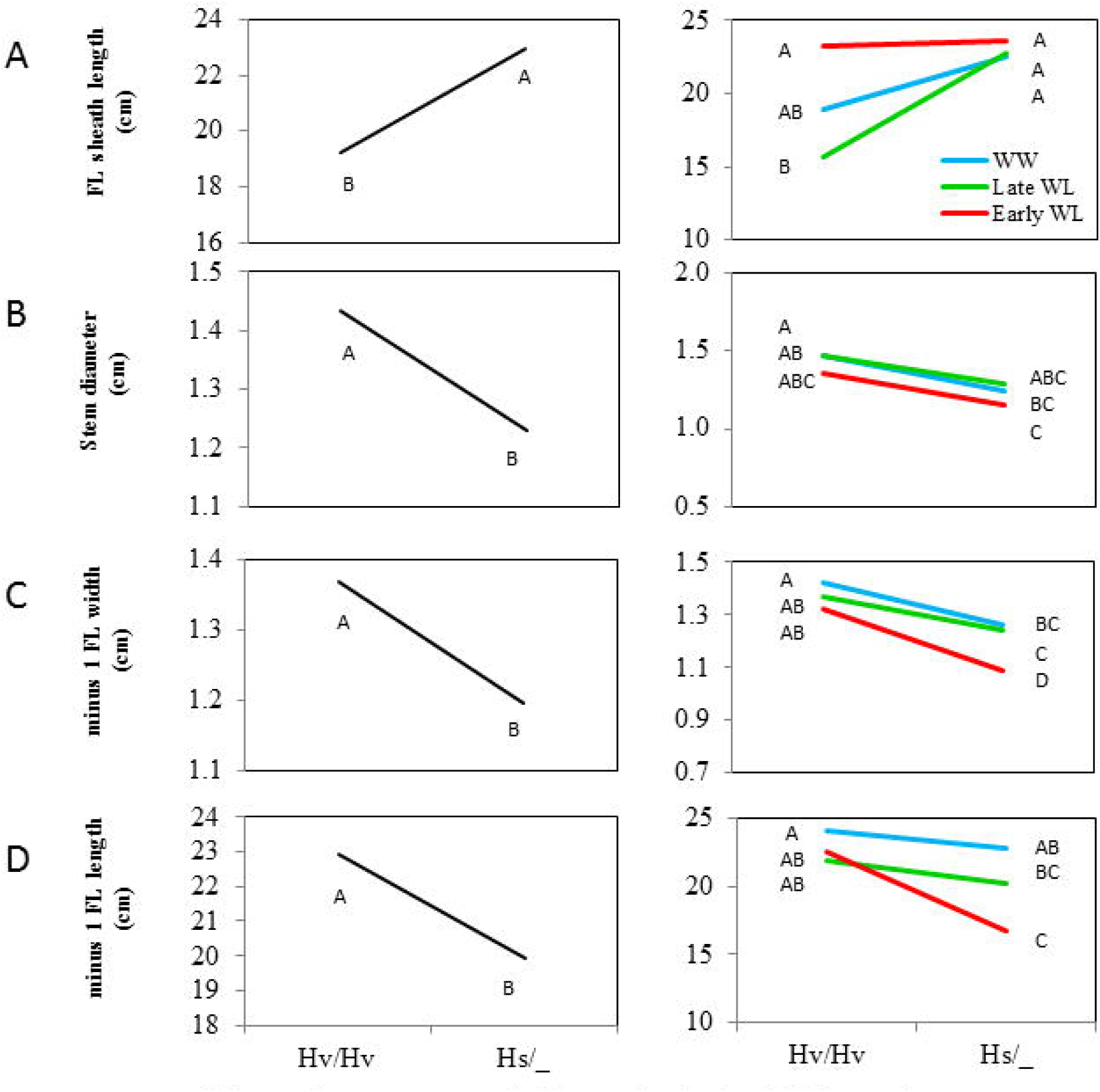
Genotype and genotype by environment interaction curves for canopy structure traits: (A) Flag leaf sheath (cm) length, (B) stem diameter (cm), (C)-1 flag leaf blade width (cm), (D)-1 flag leaf blade length. Hv/Hv, homozygous for the Barke cultivated allele, Hs/_, homozygous or heterozygous for the wild HID062 allele.

#### Leaf senescence

Analysis of senescence carried out by calculating the difference of the yellow-brown pigments area out of total canopy area during a period of 12 days (66 and 78DAP, Fig. 9A-B). No significant differences were found in total canopy area between genotypic groups in both dates, nor in yellow-brown/total at 66 DAP (not presented). However, at 78 DAP the carriers of the cultivated allele exhibited on average a significant higher yellow-brown out of total as compare to the Hs/_ group, and greater senescence, evaluated by calculating the change in the percentage of yellow-brown out of total during these 12 days, was observed for the former (Fig. 9C). Notably, these differences in senescence between the two genotypic groups were more pronounced under both drought stress (Fig. 9C).

**Fig. 9.**
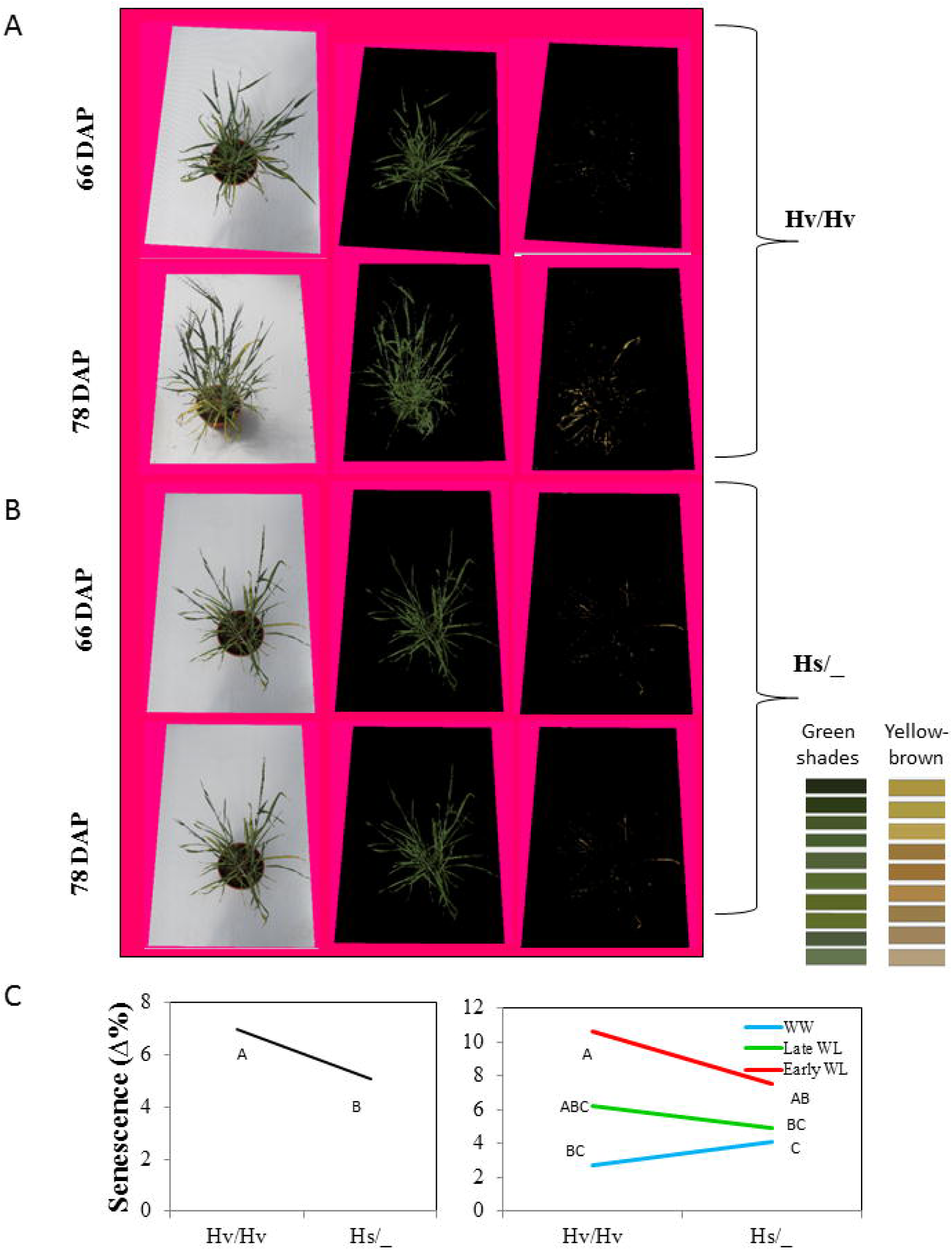
The HsDry2.2 is associated with reduced leaf senescence under drought (A) Representative plant images used for analysis of senescence (Early WL), as the difference of the yellow-brown pigments out of total canopy area between two sets of photos taken at 66 and 78DAP. Analysis was conducted using a designate home-designated software (see Methods). (B) Genotype and genotype by environment interaction curves for leaf senescence. Hv/Hv, homozygous for the Barke cultivated allele, Hs/_, homozygous or heterozygous for the wild HID062 allele.

## Discussion

### The HEB-25 genetic architecture for flowering time

The genetic architecture identified in our experiments for flowering time was similar to that reported earlier by Maurer et al. (2015) and Saade et al. (2016), with the exception of Vrn-H2 that was not identified in our study. The main loci for HEA, including HvELF3, Ppd-H1, HvCEN, *denso*, Vrn-H1, and Vrn-H3, were identified (Figs. 2 and Table S4). Nevertheless, an interesting difference between the experiment in Israel (E34°47′, N31°54′) and that conducted in Germany (E11°58′, N51°29′) was the effect of Ppd-H1 on HEA. Unlike the earliness effect of the wild allele seen in Germany, i.e. a mean 9.5-day reduction in heading time (Maurer et al. 2015), under our conditions, an opposite effect was obtained, with a 6.7-day increase in mean heading time under both growth conditions. This effect is similar to that reported by Saade et al. (2016), and highlights the role of this gene in photoperiod sensitivity (Turner et al. 2005). These type of opposite effects should be considered in “designing” an ideotype or pyramiding wild QTL to achieve these in breeding. These type of QTL originated from wild alleles could be beneficial in one environment and detrimental in other.

### Genome scan for QxE interactions

In this study, a genome scan was performed in search for loci with significant QxE interactions, as manifested by environmentally-conditioned differences in mean trait measures between the carriers of the wild allele (Hs/Hs) as compared to plants homozygous for the cultivated allele (Hv/Hv). This approach is different from recent QxE analysis in Arabidopsis, in which analysis was performed in two stages: initially performing a genome scan for loci that are associated with the trait variation per-se regardless of the environment, mainly to maximize QTL identification, and only then the QxE interactions for these loci was tested (El-Soda et al. 2014). In addition, it is different from the way GxE is often treated as a variation component that would improve the percentage variation explained by the genetic model (Elias et al. 2016). HsDry2.2 is an excellent example of a major QxE locus (Fig. 3) for reproductive output traits (PGY and GN) that was not identified by GWAS for the trait per-se in neither WW nor WL conditions (Fig. 2, Table S4-5). Yet, when the genome scan was screened for QxE, it was highlighted as the most reproducible QxE locus with similar effects observed over two years of field trials.

### Prevalence of QxE QTL for reproductive rather than vegetative or phonological traits

In this study we were able to identify 69 loci for trait per-se and 22 for QxE (Fig. 2; Table S4 and S5 respectively). In general, comparison of the QxE genomic architecture to that of the traits variation per-se shows that there are very few QxE loci for some of the traits. Moreover, some bias for reproductive trait-associated QxE loci was apparent (Supplementary Tables S4 and S5). This does not seem to be the result of the limited variation of vegetative or phonological traits in the HEB-25 lines under our experimental set-up (Fig. 1A). Moreover, we were able to identify 69 loci for trait per-se and 22 for QxE (Fig. 2; Table S4 and S5 respectively).

Interestingly, the prevalence for conditional QTL is by far more prevalent for PGY or GN as compared to other traits in which the large number of QTL for trait per se was dramatically reduced while considering interaction. Looking more carefully on the reaction norms of such QTL (Fig. 4 and Fig. S3) show that the carrier of the wild allele experienced less reduction under WL rather than simply increasing the PGY or GN i.e. the wild allele is conferring phenotypic stability under changes in the watering regime. Although this is shown for a semi-controlled environment and requires validation under a denser agricultural set-up, such locus has a promising potential to provide grain yield stability against drought. Future experiments should therefore test isogenic lines for this wild allele in several genetic backgrounds and with larger plots. It is argued that since drought survival is a trait under strong evolutionary pressure, many drought survival loci would be expected to impart tolerance in crop plants by accumulation of small effect QTL (Mickelbart et al. 2015). However, when considering grain yield and exploitation of interspecific crosses, such as the HEB-25, one caveat should be considered. Wild relatives of modern crops adapt to drought, not necessarily by producing more grains under stress, a strategy that might be detrimental for maintenance of wild populations under limited and fluctuating resources. Instead, upon dissection of the wild genome and examining its parts in a cultivated genetic background, as in our study, we would not necessarily expect to find alleles from the wild that would increase grain number and yield under stress. Rather, the alleles that we expect to find are such that stabilize or buffer the effects of the stressful environment on the plant. For example, owing to the wide allelic variation existing in the HEB-25, we were able to identify several significant QxE loci, including a major locus that implicates a hitherto unknown role for a “flowering time gene” in regulating drought resilience in what seems as a non-escape mechanism. The pot experiment showed similar trend of shortening time to flowering (only ~2.5days) of the wild allele as compare to the cultivated one. The much earlier general heading date in the pot experiment (~65days), as compare to the field trails (~100days), is probably related to differences in thermal degrees days between experiments (Fig S6). Hence, in the relatively hot climate of the pot experiment HEA was reduced for both alleles, therefore maintaining the differences between them yet to a lesser extent in the pots than in the field. Furthermore, under these hot conditions both genotypes flowered early, ruling out drought escape mechanism of one allele compared to the other. Alternatively, the earlier flowering and elongated grain filling of the wild allele may in part underlie its improved productivity.

### Flowering-independent QxE effects of HsDry2.2 on reproductive traits

The most reproducible and significant QxE locus was identified on the long arm of chromosome 2H (Fig. 2,3). Interestingly, this position matches the location of the HvCEN, the barley ortholog of the CENTRORADIALLIS gene, with two main haplotypes differentially distributed over spring and winter barley varieties (Comadran et al., 2012). Under WL conditions, HsDry2.2 showed conditional effects on the total plant grain number, but no such interaction with flowering time (Fig. 4).

Both field trails and pot experiment shows that the wild allele had no advantage over the cultivated one in vegetative or TDM production, yet, it shows superiority in terms of reproductively (PGY and HI). This phenotype presented is most probably not related to gibberellin insensitivity mechanism, as plant height of wild allele carriers was not modified compare to the cultivated allele. Most recently loci regulating developmental characteristics were found to be co-located with flowering time gene including the HvCEN (Maurer et al., 2016 and Nice et al., 2017). Yet, this locus was not associated with height differences in this study either (Nice et al., 2017). Maurer et al. (2016) found significant effects of 'QTL-2H-7' (with HvCEN as candidate gene) on all measured traits, however they did not measure yield. It would therefore be interesting to further characterize these differences at the metabolic and developmental levels, both in the sink and source tissues, under gradients of water availability for isogenic lines for this QxE locus. This may lead to the identification of hitherto unknown pathways related to drought tolerance.

### HsDry2.2 effects under early vs. late drought

Despite the similarity in PGY production for the Early and Late WL treatments, it is quite clear than the reduction was obtained via different pathways. As expected, while spike/plant was significantly reduced by the Early WL, in the Late treatment it did not. GW (all) under the Late WL was reduced half of that of the Early WL, and GN appears to be reduced only in the Early WL, although not significantly (Table S6).

As temperatures in the greenhouse were high along the entire season (Fig S6), it is reasonable to assume that all treatments plants were under mild heat stress. In addition, as plant water demand grew towards the middle of the season, the field water capacity dropped beneath 60% in the WW when irrigating once in two days (and not daily). Therefore, the advantage in PGY for the wild allele, which was obtained mainly under WW might reflect also heat resistance.

Whereas under the WW treatment the benefit of the wild allele in PGY may be attributed to higher GN, under both WL treatments the wild allele presented an advantage in GW, ripening period, and reduced senescence rate. In that respect, the condensed canopy structure of the drought adapted wild allele under Early WL may also have an effect on reduced senescence rate as a 'by product' of this phenotype.

Overall, the Hs_ plants presented a phenotype that matches the design of future climate-resilient barley ideotypes for Mediterranean climatic zones based on crop models (Tao et al. 2017), i.e. longer reproductive growing period (similar to longer RIP), lower leaf senescence rate, and higher drought tolerance. This matching indicate the importance of this QTL for future breeding of barley.

Notably, resequencing the hitherto known SNP between spring and winter barley in the segregating BC_2_S_1_ plants show that both Hs and Hv alleles (Fig. S7, Table S1) are in fact carrying G at this position termed Pro135A by (Comardan et al. 2012). These results may indicate that the casual variation underling the improved drought resistance of Hs over Hv allele in this study may be attributed to other variation than this SNP, i.e. that there is another allele that cause these effects. Possibly, this could also be other gene rather than the HvCEN itself, which is locked with this chromosomal region as was shown recently (Mascher et al. 2017), and therefore dissecting the true causal variation in this region is still challenging

Overall, the currents study goes from large-scale genome-wide association scan to a finer resolution scale, shedding some more light over the promising HsDry2.2 locus and its mode of action. This seems like a rare case of wild allele with direct beneficial effects on grain yield. Finer mapping of this locus should validate if indeed the causal gene for these multiple differences between wild and cultivated allele correspond to the CEN or nearby overlooked gene/s.

## Supplementary data

**Table S1.** Primers used for HRM and HvCEN sequencing

**Table S2.** Field distribution of traits values under WW and WL

**Table S3.** Field pairwise correlations between traits under WW and WL

**Table S4.** GWAS results for trait per se.

**Table S5.** QxE loci for the different traits

**Table S6.** Pot experiment ANOVA

**Fig. S1.** Genotyping BC_2_F_1_ pot experiment

**Fig. S2.** Irrigation management

**Fig. S3.** Field VDW reaction norms

**Fig. S4.** Field family reaction norms

**Fig. S5.** Pot experiment shoot morpology

**Fig. S6.** Pot experiment temperatures

**Fig. S7.** HvCEN sequencing

### Acknowledgements

We wish to thank Yoske Gotlib and the staff of the Experimental Farm at the Faculty of Agriculture, Rehovot), Matan Koren, Khaled Bishara, Oshreet Finer and Ira Bartkiv (ARO, Bet Dagan) for their technical assistance. The discussions and technical assistance of Prof. Menachem Moshelion (The Hebrew University) were instrumental for setting the high-content phenotyping system. We also thank Yehudit Posen for editing the manuscript. The ERA CAP Grant Pi339/8-1 to K.P and E.F supported this research. The authors declare that they have no conflict of interest

## Contributions

R.S., L.O.M, E.L and E.F. designed the experiments, collected the data, made the data analysis and interpretations and wrote the manuscript. A.F. and L.O.M. were involved in the data analysis, its interpretations and assisted in writing the manuscript. K.P. and A.M. supplied the plant material and genotype data.

## Figures legends

**Fig. S1** (A) High resolution melting analysis for the peak marker BOPA2_12_30265 for the genotyping of BC_2_F_2_ (self of backcrossing line HEB-04-96) segregating population in the HsDry2.2 locus. (B) Sanger sequencing showing the different alleles for the BOPA2_12_30265 SNP.

**Fig. S2** The effects of water limitation on control plants grown in a semi-controlled, high content plant phenotyping (HCPP) set-up. (A) The experimental set up for the 2015 and 2016 experiments included the HEB-25 lines and control plants. Plants were grown in pairs of troughs, under well-watered (WW) or water-limited (WL) conditions. (B) Accumulated irrigation and (C) stomatal conductance as a function of days from sowing, under the WW (blue) or WL (red) conditions, during 2016. (D) A panoramic view over the pot experiment at ripening stage. (E) Field capacity (%) as a function of days after planting (DAP) for the different genotypes. Seedlings were planted into small pots (60g soil and 300g water, 100% field capacity), at 28 DAP plants were transplanted into medium pots (190g soil and 1170g water), ant at 52 DAP plants were into large pots (460g soil and 2500g water). The Early WL initiated at planting and the Late WL at 52 DAP, about two weeks before booting. Pots were weighed manually before and after irrigation, keeping the well-watered (WW) pots between 60-90% of field capacity and the water limited pots (WL) at 40-60%. (F) Stomatal conductance measured for cv. Barke control at 56 and 69 DAP under the different irrigation treatments.

**Fig. S3** Reaction norm of HsDry2.2 is illustrating the mean values of vegetative dry matter (VDW) under WW and WL conditions, in the whole HEB-25 population. The three genotypic groups of plants homozygous for the Barke cultivated allele (Hv/Hv), homozygous for the wild allele (Hs/Hs) and heterozygous (Hv/Hs) are depicted by blue, red and gray lines, respectively.

**Fig. S4** Reaction norms of HsDry2.2 are illustrating the mean values of plant grain number (GN) under WW and WL (reaction norms) conditions, in the a) HEB-16 and b) HEB-05 families. Homozygous for the Barke cultivated allele (Hv/Hv) and for the wild allele (Hs/Hs) are depicted by blue and gray lines, respectively. Heterozygous plants were excluded from analysis due to low number of replicates (<6).

**Fig. S5** (A) Representative photos for canopy structure modifications, showing the carriers of the wild allele (Hs/_) have on average longer sheath, and narrower-shorter leaf blades (of FL and −1FL). (B) Stem cuts after harvest at the base of the spike (Late WL) showing Hs/_ plants to have reduced stem diameter, which also appear to be thicker than in the cultivated allele.

**Fig. S6** Air minimum and maximum daily temperatures monitored over the pot experiment season.

**Fig. S6** Sanger sequencing results for the known SNP between spring and winter barley in the segregating BC2S1 plants. Both Hs and Hv alleles are in carrying G at this position termed Pro135A by Comardan et al. (2012) whereas additional cv Barke control plants experiment are shown to carry C.

**Table S1.** Distribution and heritabilities of traits values under WW and WL. Wide-sense heritability is calculated by ANOVA as the proportion of the phenotypic variation explained by the genotype (family) effect in a multi-factorial model.

**Table S2.** Pairwise correlations between traits under WW and WL

**Table S3.** GWAS results for trait per se. The effect correspond to the percent difference between mean phenotypic value of homozygous for the wild allele compare to carriers of the cultivated Barke allele within the whole HEB-25 population

**Table S4.** GWAF results for QxE. The effect is calculated as the difference between the effect of the wild allele under WW and WL. Positive values indicate higher increasing or less reducing effect of the wild allele under WL.

**Table S5.** Least square means of the measured traits: total DM (TDM, gr/plant), plant grain yield (PGY, gr/plant), harvest index (HI), spike per plant, grain number of three first spike (GN-first three), grain number (GN all), grain weight of three first spike (GW-first three), grain weight (GW, all), days from planting to booting (HEA, days), ripening period (RIP, days), Flag leaf sheath (cm) length, stem diameter (cm), −1 flag leaf blade width (cm), −1 flag leaf blade length.

**Table S6.** Analysis of variance (ANOVA) for the measured traits: total DM (TDM, gr/plant), plant grain yield (PGY, gr/plant), harvest index (HI), spike per plant, grain number of three first spike (GN-first three), grain number (GN all), grain weight of three first spike (GW-first three), grain weight (GW, all), days from planting to booting (HEA, days), ripening period (RIP, days), Flag leaf sheath (cm) length, stem diameter (cm), −1 flag leaf blade width (cm), −1 flag leaf blade length.

